# Dynamic representation of space in the hippocampus: spatial novelty detection and consolidation in CA1 and CA2

**DOI:** 10.1101/2021.08.19.456964

**Authors:** Guncha Bhasin

**Affiliations:** Division of Systems Neuroscience, National Brain Research Centre (NBRC), Manesar, Haryana, 122051, India

## Abstract

Hippocampal place cells are the functional units of spatial navigation and are present in all subregions-CA1, CA2, CA3 and CA4. Recent studies on CA2 have indicated its role in social and contextual memory, but its contribution towards spatial novelty detection and consolidation remains largely unknown. The current study aims to uncover how CA1 and CA2 detect, process, assimilate and consolidate spatial novelty. Accordingly, a novel 3-day paradigm was designed where the animal was introduced to a completely new environment on the first day and to varying degrees of familiarity and novelty on subsequent days, as the track was extended in length and modified in shape, keeping other environmental constraints fixed. Detection of spatial novelty was found to be a dynamic and complex phenomenon, characterized by different responses from hippocampal place cells, depending on when novelty was introduced. Therefore, the study concludes that early novelty detection (the first time a novel space is introduced in a relatively familiar environment) and subsequent novelty detection are not processed in the same way. Additionally, while neuronal responses to spatial novelty detection (early and subsequent) were found to be the same in CA1 and CA2 ensembles, their responses differed in spatial consolidation mechanisms during subsequent sleep replays. For CA1, spatial coverage of prior behaviour was found to be closely reflected in subsequent sleep for that particular day, but CA2 showed no such ‘coherent response’, highlighting mnemonic processing differences between CA2 and CA1 with respect to spatial novelty.

## Introduction

The discovery of place cells in the hippocampus more than a quarter of a century ago (O’Keefe & Dostrovsky, 1971) firmly establishes this region as the locus of the cognitive map in the brain, essential for successful spatial memory and navigation. Subregions of the hippocampus such as CA1, CA3 and DG have been extensively studied and their respective roles characterized conclusively, (Wilson & McNaughton, 1993; Zhang *et al*., 1998; Leutgeb *et al*., 2007; Mizuseki *et al*., 2012; Kim *et al*., 2020) but CA2 has largely been left out of the spotlight, and termed as a mere ‘transition zone’ with no functional contribution of its own towards this map (Ishizuka *et al*., 1995) . This has since been rectified with many studies in the last decade, which have focused on structural, functional and unique connectivity patterns of CA2 and it being a key player in the hippocampal circuitry, highlighting its important contribution towards memory processing (Kohara *et al*., 2014; Stevenson & Caldwell, 2014; Mankin *et al*., 2015; Dudek *et al*., 2016; Kay *et al*., 2016).

A key role of spatial exploration is successfully detecting and encoding novel stimuli (contextual or social) and integrating it with familiar stable spatial maps of an environment, effectuating continual upgrading of stored memories. In previous studies, CA1 has been recognized as a broadcaster of a novelty signal in the hippocampus (Larkin *et al*., 2014) and disruption of EC-CA1 pathway leads to impairment in spatial novelty detection (Vago & Kesner, 2008). With the discovery of the novel disynaptic pathway (EC(II) to CA2 to CA1) (Chevaleyre and Siegelbaum,2010 ; Kohara *et al*,2014) which is parallel and independent to the ‘classic’ trisynaptic pathway (EC to DG to CA3 to CA1), CA1 is uniquely poised to be neuromodulated through diverse circuitries within and beyond the hippocampus by both CA3 and CA2 (Mankin *et al*., 2015). While CA3-CA1 pathway has been extensively studied with respect to spatial navigation and memory, with some even looking at CA3-CA2 place cell topography (Lee *et al*., 2015; Lu *et al*., 2015) CA2-CA1 studies are far and few in between (Mankin *et al*., 2015; Alexander *et al*., 2016). Additionally, the mutual inhibitory relationship between CA3-CA2, controlled by feed-forward circuitry (Chevaleyre & Siegelbaum, 2010; Kohara *et al*., 2014) and limited plasticity (Zhao *et al*., 2007) indicates a competition for active control of the hippocampal circuitry. All these aforementioned studies establish that multiple independent circuitries co-exist within the hippocampus, which are activated simultaneously or alternately for different functional optimization; with maximum influence being exerted on the most downstream subfield of the hippocampus, i.e. CA1. Recent studies have shown that CA2 plays a role in social and novel contextual information processing (Hitti & Siegelbaum, 2014; Mankin *et al*., 2015; Alexander *et al*., 2016) and has a relatively flexible spatial code when compared to that of CA1 and CA3. Furthermore, axons of CA2 basket cells extend to all 3 CA fields of the hippocampus, providing feedback inhibition to CA3 and feed-forward inhibition to CA1, making it uniquely poised to co-ordinate and control the hippocampal circuitry (Mercer *et al*., 2007).

Place cell activity during prior awake behaviour is reflected in ‘replays’ during sharp wave ripples (SWRs) that occur during nonrapid eye movement (nREM) sleep (Kudrimoti *et al*., 1999; Nádasdy *et al*., 1999; Lee & Wilson, 2002). During these SWRs, place cell ensemble activity is replayed at faster timescales and spatial topography of familiar and novel space is encoded, (Nádasdy *et al*., 1999; Brun *et al*., 2008; Dragoi & Tonegawa, 2013; Wu & Foster, 2014) indicating that sleep is critical to memory consolidation of previous experiences. While orderly time-compressed replays primarily reflect trajectories navigated in prior behaviour, the spatial content of many SWR ripples remains unknown (Chen *et al*., 2016). Therefore, understanding the firing patterns and frequency of firing of hippocampal place cell ensembles during sleep across all replay events, beyond their temporal order and directionality (forward/reverse replays) is imperative to a better understanding to memory consolidation mechanisms and spatial information processing of novel and familiar topographies in sleep.

The primary goal of the present study is to tease out the specific contributions of CA1 and CA2 towards mnemonic processes such as detection of novel spatial stimuli and it’s subsequent consolidation. To meet this end, in vivo multiunit recordings were performed in rat hippocampus to examine and distinguish between neuronal firing responses in CA1 and CA2 to spatial novelty detection and consolidation. A novel 3-day paradigm, where the rat experienced varying degrees of spatial familiarity and novelty with each successive day, as the track was elongated and geometrically reshaped, was specifically designed for this aim. Across days (days1 v/s day2 v/s day3) and within days comparison (days 2 and 3: novel v/s familiar) between CA1 and CA2 ensemble populations were made with respect to place cell counts, average firing rates and pairwise cross correlations between cell pairs, separately for place cells that fired on the elongated portion of the tracks on day2 and 3 and for the common portion of the track from day1 that was present on all days. Furthermore, spike activity from CA1 and CA2 place cells within sharp wave ripples (SWRs) in sleep during replay events from each day were analyzed to understand how the consolidation of an ever changing track/environment as it gets modulated in shape, form and size occurs across days. As an outcome, the ensemble activities of neural populations in CA1 and CA2 were estimated and interpreted to tease out the role of CA1-CA2 system in various aspects of novelty detection and consolidation across days during a dynamically evolving spatial navigation task.

## Materials and methods

### Animal handling and surgical procedures

5-6 month old Long-Evans rats (n = 4, male) were housed individually on reversed light-dark (12:12 h) cycle and the experiments were carried out during the dark phase of the cycle. All surgical procedures were performed under aseptic conditions. The rats were initially anaesthetized using ketamine (administered at 60 mg/kg b.w.) and xylazine (administered at 8mg/kg b.w.) and subsequently shifted to gaseous anaesthesia using isofluorane for the rest of the surgery. A custom-built recording device (Microdrive) contained inside a dual bundle, was made entirely from scratch in the laboratory, containing 20 independently movable tetrodes (each bundle consisting of 9 recording tetrodes +1 reference tetrode). This drive was then surgically implanted over the right hemisphere at 3.5-3.7 mm posterior to bregma and 1.7-1.8 mm lateral to midline; to simultaneously access different regions of the hippocampus (CA1 and CA2). (fig1B).

**FIGURE1:**
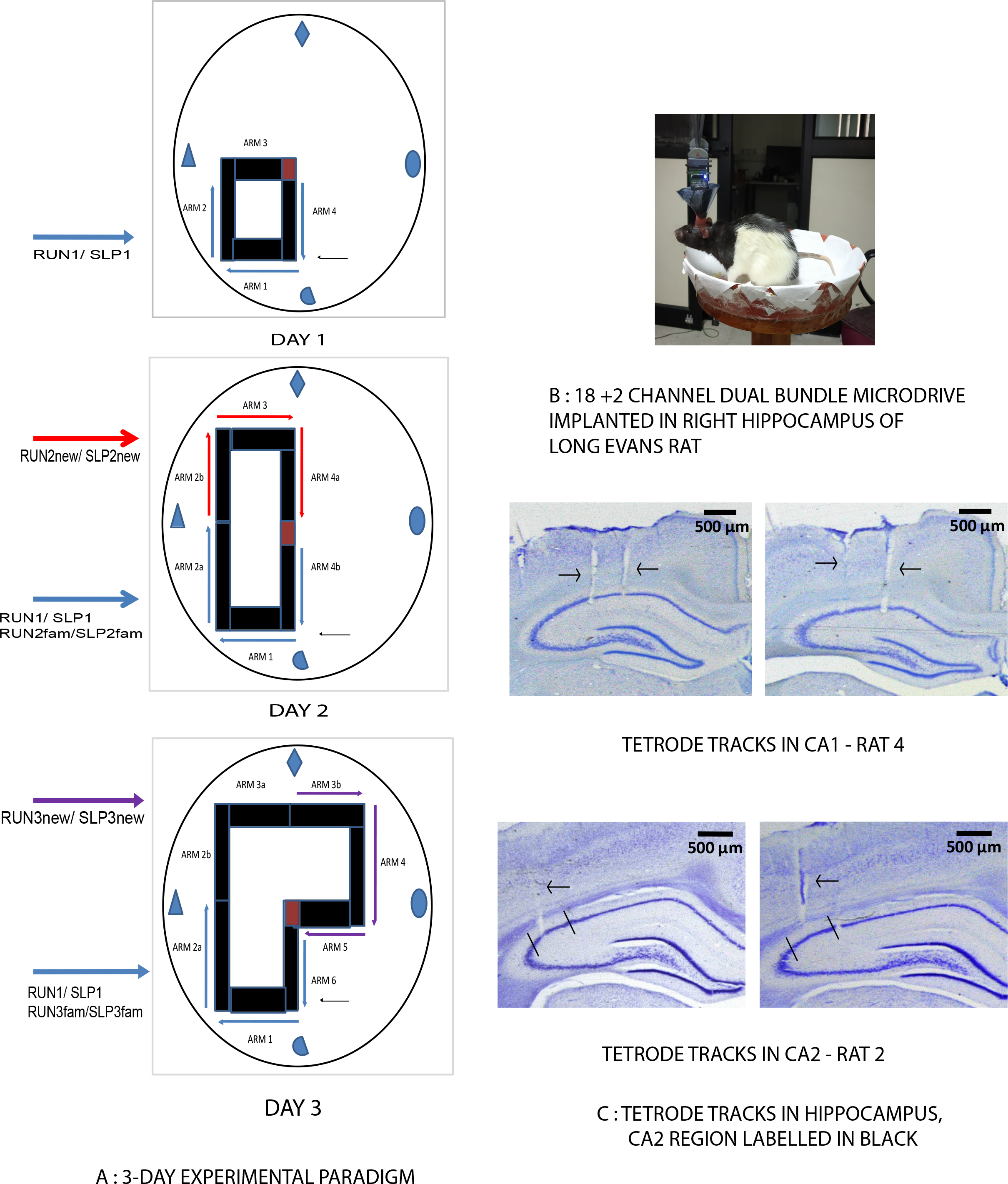
a) Schematic of the experimental tracks used in the novel 3-day novel paradigm: on day1, a square track of 4 equal arms was placed in 1 corner of the room. Arms 1,2,4 are termed as RUN1 and are a part of all 3 tracks. On day2, the previous track was extended to double its size. The extended arms are termed as RUN2new, while RUN1 becomes RUN2fam. On the 3^rd^ day, the track was extended by the same size C-shaped track used on day2, termed RUN3new, and RUN1 becomes RUN3fam. The middle arms (2b and 3a) are not used for within day comparisons to maintain consistency of sizes of track used for comparison. The placement of reward corner (yellow corner), along with 4 perpendicularly placed distal visual cues on the wall is consistent across all tracks on all days. b) A dual bundle 20 channel microdrive (9 tetrodes + 1 reference channels in each bundle) impanted in right hemisphere of Long evans rat, aimed at CA1 and CA2 subfields of the hippocampus. c) Tetrode tracks in CA1 (top) and CA2 (bottom) in 40 micron thick, nissil-stained histological slices of the rat hippocampus (coronal slices). CA2 region is marked with black outlines in the bottom slices (Kohara *et al*., 2014).

### Post surgical procedures

The rats were given a post-surgery recovery period of 7 days, where post-surgical care was provided by the experimenter. Following post-surgical recovery, the tetrodes were slowly advanced, targeting CA1 and CA2 regions of the hippocampus over a period of 10-15 days, by keeping the rat on a pedestal next to the recording system. During this period, the rats were also trained in the adjacent behavioural room to run clockwise, seeking chocolate sprinkles placed at random locations on a centrally placed black circular track (10 cms wide, elevated 90 cm from floor level) for 30 min/day for 8-10 days. The behaviour room was covered with circular plain black curtains at its perimeter and had no other cues (proximal or distal) of any kind. During training and subsequent experimental recordings, the rats were maintained at 85% of their free feeding weights.

### Electrophysiology and recording

17 μm platinum-iridium wire from California Fine Wire, USA was used to make the tetrodes and the tips of individual wires of these tetrodes were electroplated with platinum black solution (Neuralynx Inc., USA) to 100-150 kΩ with 0.2 μA current. Multi tetrode electrophysiological recordings were carried out using 96-channel data acquisition system (Digital Lynx 10S, Neuralynx Inc., USA) by amplifying the signals through a headstage preamplifier (Neuralynx Inc., USA). The microdrive was fitted with a EIB-27 board at it’s centre, which then connected to the headstage preamplifier HS-27, which was in turn connected to the commutator using HS-27 tethers. The commutator was connected with the recording cables to the 96-channel Digital 145 Lynx 10S data acquisition system (Neuralynx Inc., USA). The units were recorded against a reference electrode from that particular bundle of the dual – bindle microdrive, which was present in a cell-free zone in the brain (the ‘silent zone’ above the hippocampal pyramidal layer) by filtering the signal between 600 Hz and 6 kHz. Spike waveforms above the threshold were sampled at 32 kHz for 1 ms. Local field potentials (LFPs) were recorded against a ground screw anchored to the skull above the frontal cortex, filtered between 0.1 Hz and 1 kHz, and continuously sampled at 4 kHz. (fig2B). The position and the head direction of the animal were tracked with the red and green LEDs attached to the headstages, which was captured through a color CCD camera that was mounted at the ceiling, central in position to the room (CV-S3200, JAI Inc, San Jose, USA) at 25 Hz.

**FIGURE2:**
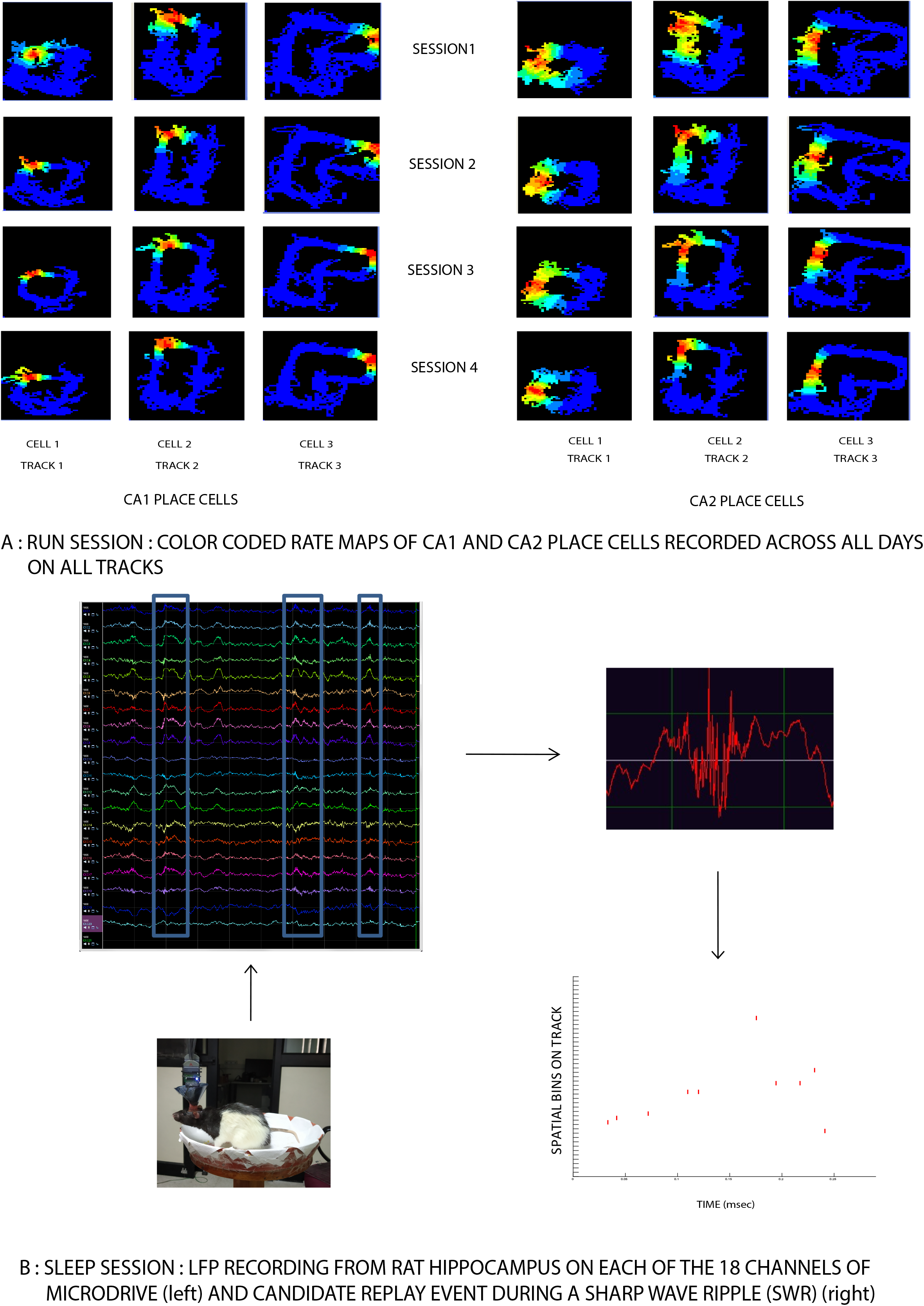
a) Examples of CA1 and CA2 place cells recorded in RUN session across all days on all 3 tracks. These are color coded firing rate maps plotted for each day’s recording, which consisted of 4 sessions of 20 clockwise laps each. b) LFP recordings from the hippocampus on one channel from each of the 18 tetrodes of the microdrive. The 2 reference channels are silent. Sharp wave ripples (SWRs), the signature of hippocampal LFP are highlighted in blue (left panel). A candidate replay event (forward replay) during one such SWR is shown in the right panel.

### Histological procedures and identification of recording sites

After successful completion of the electrophysiological experiments, marker lesions were performed on few selected tetrodes by passing current at 10 μA for 10 seconds. Rats were transcardially perfused the next day with 4% formalin solution, the brain was extracted and stored in 30% sucrose-formalin until it sank in the solution. These brains were then sectioned in the coronal plane (40 μm thick), mounted, and stained with Nissil’s staining using 0.1% Cresyl violet. Images of serial sections were captured on Leica DFC265 digital camera attached to Leica M165-C stereo microscope and saved as TIFF files. The distance from midline to the tetrode track markings were measured from these serial sections, plotted in excel spreadsheet to visualize the configuration of tetrode tracks. The tetrodes were identified by comparing this configuration with the arrangement of tetrodes in the microdrive, and cross verifying with the marker lesions. Depth reconstruction of the tetrode tracks was performed to identify the brain region at which the cells were recorded on each day, based on the distance from the bottom tip of the tetrode by taking into account 15% shrinkage of tissue due to histological processing (fig 1C).

### Experimental procedures

At the beginning of the experiment, 4 distinct shaped visual cues were hung over the black curtains at 90 degrees to each other. A large 1X1 foot black square platform was kept in the centre of the room, elevated 90cms from the ground, over which all the differently shaped, closed loop plain black tracks (10cms wide) were subsequently introduced each day, all elevated at 15 cms from the square platform. On day 1, a small square black track (45X45 cms) was kept on the lower left quadrant of the square platform. This track was divided into 4 equal length arms: arm1, arm2, arm3, arm4. The corner junction of arm3 and 4 was covered with yellow sandpaper (10cmsX10cms), which served as the location of reward for the rat (a single chocolate sprinkle) each time it completed a lap. The position of this corner was at the centre of the behaviour room, and remained constant throughout the length of the experimental paradigm. (fig1A-day1). On day2, the square track from day1 was elongated into a rectangle shaped black track (45X80cms), such that the track now occupied the lower and upper left side of the square platform (2 quadrants). The track was further divided into 6 arms: arms1,2a and 4b were the same arms from day1 track, while arms 2b, 3 and 4a were the additional/novel arms on day2. The reward corner, between arms 4a and 4b, was at the same physical location i.e. centre of the room (fig1A-day2). On the 3^rd^ day, the rectangular track from day 2 was elongated to extend to the top right quadrant of the platform and formed a L-shaped track (3 quadrants). This track was divided into 8 arms: arms 3b,4 and 5 were the additional/novel arms, arms 1,2a and 6 were the same arms from day1 track and arms 2b and 3a were the 2 remaining arms from the novel arms of day2. The position of reward area, now at the junction of arms 5 and 6 was again at the same location as previously mentioned (fig1A-day3).

All days had 4 sessions of run (fig 2A), consisting of 20 clockwise laps each (except day 1 of 1 rat, which had 3 sessions). This novel paradigm was designed such that at each elongation of the track, from day 1 to day 2 to day 3, the rat took a longer route to complete each lap. This was done so that the rat experiences a completely novel environment on day 1 (0%familiarity, 100% novelty) and varying degrees of familiarity and novelty on day 2 (50% familiarity, 50% novelty) and day 3 (67% familiarity and 33% novelty) of track environment. Except this change in the length and thus, the shape of the environment, the relative position of distal visual cues, the position of reward corner, the entry point of rat on track (arm1) and the direction of running (clockwise) all remained constant throughout the experiment. Once the tetrodes reached the hippocampal layer, their position were not disturbed or moved by the experimenter on any day of the experiment. Each session was interleaved with 10-15 seconds of break where the rat was removed from track and put in a box to dispense its sense of direction and orientation. Then it was released on the same starting position on the track for the next session. On completion of all the experimental sessions, the track was wiped clean with 70% ethanol to clear off any traces that could act as potential cues for next day of recording.

The experiment was further divided into 2 stages: RUN session, where the animal ran 4X20 laps, followed by post run SLEEP session for 3-4 hours.

#### RUN SESSION

On day 1, although the rat ran in a clockwise fashion on a square track, only place cells from arm1, 2 and 4 were considered for appropriate comparison and analysis. This is termed RUN1. Since these arms are present on both day 2 (arms1, 2a and 4b) and 3 (arms1, 2a and 6) as well, though numbered differently to maintain arm count continuity each day, they are the only common arms between each track, across all days. They are termed RUN2fam and RUN3fam respectively on day 2 and day 3. Correspondingly, the added/novel arms on day 2 (arms 2b, 3 and 4a) and day 3 (arms 3b, 4 and 5) are termed RUN2new and RUN3new, respectively. All the above mentioned C-shaped, 3-arm tracks: RUN1, RUN2fam, RUN2new, RUN3fam and RUN3new are similar in shape and size and are thus used for comparative analysis with each other. The middle arms (arms 2b and 3a) on day 3 are excluded from the same to maintain track length accuracy for analysis (fig1A). Therefore, not only can RUN1 be compared across days with RUN2fam and RUN3fam, as this area becomes more and more familiar from being completely novel across days; it can also be compared with RUN2new and RUN3new, since all 3 are novel areas of the track when first introduced in the environment on their respective days. Furthermore, comparison within the same day between familiarity and novelty can also be done on day 2 (RUN2fam v/s RUN2new) and day 3 (RUN3fam v/s RUN3new). The same comparisons have been applied to sleep analysis as well.

#### SLEEP SESSION

Each day’s recording was followed by the rat being placed on a pedestal in a room adjacent to the behaviour room, where he slept afterwards and 3-4 hours of sleep was recorded. The experimenter was present throughout this sleep recording and only recorded when they visually inspected that the rat had fallen asleep, with his eyes closed and there was no movement made by him for at least 5 minutes. During any intermittent waking of the rat (if any), the recording was switched off and resumed only after he fell asleep again. It was observed that on all days, usually the rat slept within 15-20 minutes of being brought outside the behaviour room and kept on the pedestal after completion of the experimental task.

### Data analysis

All quantitative analysis of data was performed with custom-written software on MATLAB (R2013a, 2018), (www.mathworks.com) as described below. The significance value (alpha) of all statistical tests performed is set at 0.05, unless specified otherwise.

### Isolation of single-units

Isolation of single-units was performed manually with custom-written spike-sorting software Winclust (Savelli *et al*., 2017). Cells were isolated based on the peak amplitude and energy of the waveforms recorded on four wires of the tetrode. Offline spike sorting of multiple clusters recorded from the same tetrode was done by principle component analysis (PCA) and isolating each cluster in various projections (six projections for four wires of each tetrode: 1/2,1/3,1/4,2/3,2/4,3/4). Based on their isolation quality (distance from the background and separation from other clusters), the units were rated on a scale ranging from 1 to 5 (1-very good; 2-good; 3-fair; 4-marginal; 5-poor) and the units rated ‘fair -marginal’ and above were used for further analysis. The same was done for clusters recorded in sleep as well. RUN clusters and SLEEP clusters recorded on the same tetrode were then compared and each isolated cluster’s corresponding boundaries were overlapped with each other across all six projections and only those sleep clusters that could be successfully identified with their corresponding run cluster (50% overlap or more in most projections) were choosen for further analysis.

### Linearization of tracks

Position data for all 4 sessions of a particular day for each rat was loaded in MATLAB, and outer and inner boundary was defined, in alignment with the track by removing outliers. The position where the rat was left on track at the beginning of each run session was defined as the starting point of arm1 and subsequent arms of each track were defined clockwise. All tracks were linearized and converted to 1 dimension tracks and divided into 2 cm bins. Finally, a speed filter of 2cm/sec was applied to the same.

### Defining place fields

Spatial information score: The spatial information score was calculated for all previously chosen clusters defined as place cells. The spatial information value indicates the amount of information about the rat’ position, conveyed by the firing of a single spike from a cell (William E. Skaggs &Bruce L. McNaughton, 1992), and was calculated as:

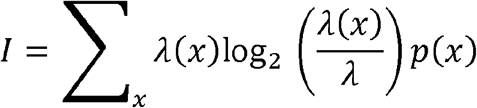

where x is spatial bin, λ(x) is the firing rate of the cell at location, λ is the mean firing rate and p(x) is the probability of occupancy at bin x. A cell was classified as a Place cell if its spatial information score was found to be significant (*p* < 0.05) based on cell shuffling procedure (defined below) performed on each cell individually in any of the experiment sessions recorded in a particular day.

Data shuffling procedure: An individual cell’s entire spike sequence recorded in a particular session was time shifted by a random interval between 20th second and 20 seconds before termination of that session. Spikes exceeding the total time of the session were wrapped around to be assigned to the beginning of session to generate new spike time sequence for that cell. This procedure was repeated 100 times for each cell separately. The 99th percentile value of the shuffled distribution of each score was taken as the threshold value for that particular cell (Langston *et al*., 2010).

The chosen clusters that fit the criteria for being defined as a place cell based on its spatial information score were subsequently loaded in MATLAB and only those clusters that fired more than 50 spikes in each session were eventually chosen for analysis. The firing rate was calculated as the ratio between the number of spikes and time spent in each bin. Linearized ratemaps of 2 cm spatial bins were smoothed with a Gaussian smoothing function of 4 cms standard deviation and characterized as having a firing rate greater than 1Hz over a minimum of 5 continuous spatial bins and an occupancy rate greater than 0.1Hz. Place fields were further defined as having a mean firing rate between 0.1Hz and 5 Hz and a peak firing rate of minimum 2 Hz. A cell’s peak rate was defined as the firing rate in the bin with the highest rate on the linear track. Place field borders were defined as where the firing rate fell to less than 10% of peak firing rate of the cell or less than 1Hz, whichever was higher. Firing rates for place cells firing in multiple sessions were averaged for the day, across all sessions. Place field centres were calculated for all place cells based on the spatial bin that had the highest firing rate for that particular cell (Dragoi & Tonegawa, 2013; Pfeiffer & Foster, 2013). Using place field centres, cells were arranged on each track for a particular day from RUN start till end in the direction the rat traversed the tracks (clockwise). Place cells firing for reward area each day were eliminated in both run and sleep and were not used for further analysis.

### Offline detection of SWRs and sleep analysis

The tetrode that had a clean LFP pattern by spectrogram (plotted in MATLAB) and showed clear SWRs (viewed in neuraview software, by neuralynx) were choosen for each day. LFP envelope was computed by first band pass filtering and subsequently taking the hilbert transforms between 140-250Hz. REM and nREM phases were distinguished by computing delta/theta ratio of 2 -3 (theta band: 6-12 Hz, delta band : 1-4Hz) using Thompsons multi taper estimate (Thomson, 1982). Within the nREM epochs, each SWR was identified when the signal exceeded 3SD (standard deviation) for 15 msec or more, and its beginning and end was marked by 1SD (standard deviation). Candidate replay events lasting >500ms were discarded. Only spikes firing within the interval of each sharp wave ripple event were considered for further analysis (Csicsvari *et al*., 2007; Karlsson & Frank, 2009).

Replay sequences were identified using an algorithm, previously described in detail in (Gupta *et al*., 2010), that detects sequence structure in the pattern of place cell activity by comparing the times and place cell centers of spike pairs occurring in a flexible time window within each identified SWR. At least 4 spikes firing from 3 different place cells within a SWR was termed as a candidate replay event. For cells with multiple place fields, spikes were assigned the place field center that maximized the forward or backward score. This algorithm resulted in a series of time windows, place field center-labeled spikes, and scores for each forward and backward sequence.

These sequences were then analyzed to identify significant sequence replays using two independent bootstrapping procedures: spike time shuffle and centre peak shuffle. The first method involved shuffling of spiketimes, while preserving the identity of spikes of each cell while the second involved shuffling of peak firing position of each cell, keeping the cell identity intact. Each event was shuffled 300 times, and its sequence score was computed again. If the unshuffled replay score was greater than 85% of both independent sets of shuffled replay scores, they were deemed to be significant and were chosen for further analysis.

## Results

A total of 427 CA1 and CA2 place cells (CA1: 288, CA2: 94, rest were cells that fired for reward area) were identified in RUN and 275 (CA1: 192, CA2: 77, rest were cells that fired for reward area) in SLEEP. After excluding the place cells that fired for the reward area from the 427 place cells mentioned above, 105 place cells were recorded on day1, 136 place cells on day 2 and 141 on day 3.

### Spatial novelty detection : early and subsequent detection

The aim of this study was to elucidate differences between CA1 and CA2’s contribution towards spatial novelty detection (early and subsequent) and consolidation mechanisms. To better understand the processing behind early and subsequent novelty detection at neuronal ensemble level, the total number of place cells were compared across days (RUN1 v/s RUN2fam v/s RUN3fam), (RUN1 v/s RUN2new v/s RUN3new) and within days (RUN2fam v/s RUN2new) and (RUN3fam v/s RUN3new), separately for CA1 and CA2.

### Place cell counts

When the track was elongated each day, from day 1 to 2 to 3, it was seen that there was an overall increase in total place cell population counts of CA1 from day 1 to day 3 and for CA2 from day 1 to day 2 with a slight decrease on day 3(fig3A). But when this comparison was made for each RUN session, it was seen that while from RUN1 to RUN2 (RUN2fam + RUN2new) cell populations increased in number, it decreased from RUN2 to RUN3 (RUN3fam + RUN3new). This was also observed for CA1 and CA2 separately, despite the fact that RUN2 and RUN3 track lengths are the same (fig3B).

**FIGURE 3:**
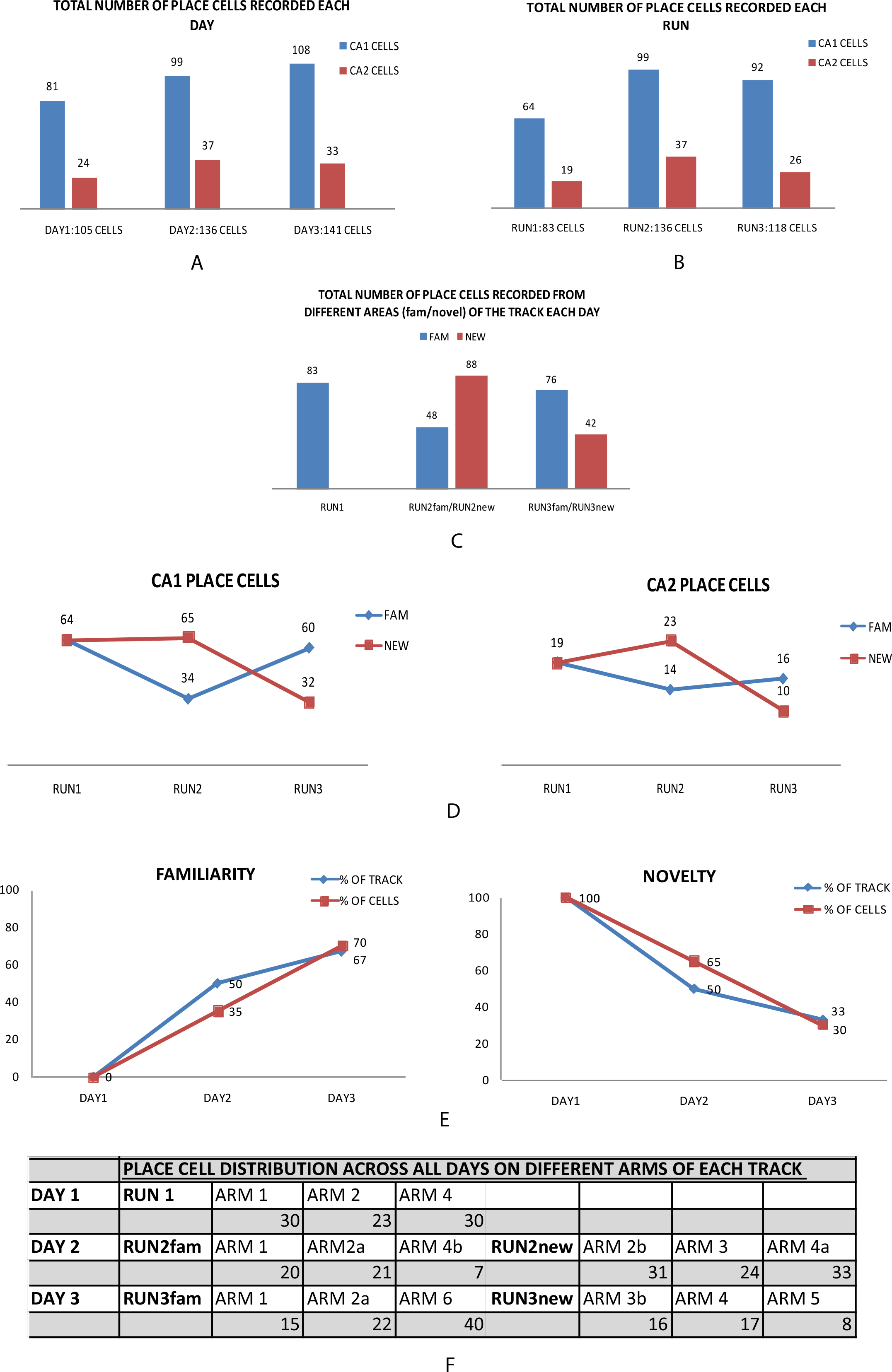
a) Total number of CA1 and CA2 place cells recorded each day (eliminating place cells that fired for reward area). b) Total number of CA1 and CA2 place cells recorded in each RUN session (RUN1:arms1,2,4; RUN2: arms1,2a,4b(RUN2fam)arms2b,3,4a(RUN2new); RUN3: arms1,2a,6(RUN3fam) arms3b,4,5(RUN3new)). c) Total number of CA1 and CA2 place cells recorded from different areas of the track (familiar and novel areas) in each RUN session (RUN2 divided into RUN2fam+RUN2new and RUN3 into RUN3fam+RUN3new). d) Comparison of CA1 and CA2 place cell distribution across RUN1, RUN2fam, RUN2new, RUN3fam, RUN3new. Both ensembles show the same trend in place cell allocation as new tracks are added on days 2 and 3. In RUN2, for both ensembles RUN2new>RUN2fam and in RUN3, RUN3fam>RUN3new for place cell distribution on tracks. 1-tailed, paired T-Test: CA1: RUN2fam/RUN2new : p=.074, RUN3fam/RUN3new: p=.007; RUN2fam/RUN3fam: p=.017, RUN2new/RUN3new:p=.056; CA2: RUN2fam/RUN2new:p=.038, RUN3fam/RUN3new:p=.006; RUN2fam/RUN3fam:p=.09, RUN2new/RUN3new:p=.02. e) As familiarity increases and novelty decreases across days (and across each RUN) on the track, place cell distribution for both CA1 and CA2 closely mirrors this trend on day 1 (100% place cells for 100% novel track) and day 3 (70% place cells for 67% familiar track and 30% place cells for 33% novel track). The same is not observed on day2 where a skewed distribution of place cells (65% place cells for 50% novel area, 35% place cells for 50% familiar area) occurs, when novelty is introduced in a relatively familiar environment for the first time (termed early novelty detection). f) Place cell distribution of both CA1 and CA2 place cells across all arms of tracks being used in comparison: RUN1, RUN2fam, RUN2new, RUN3fam, RUN3new. Place cells were found to be pretty evenly distributed across individual arms of the above tracks on all days. Kruskal Wallis test : RUN1 arms place cell distribution (arm1/arm2/arm4): p=.9, RUN2fam arms (arm1/arm2a/arm4b) :p=.06, RUN2new arms (arm2b/arm3/arm4a): p=.67; RUN3fam arms (arm1/arm2a/arm6): p=.058, RUN3new arms (arm3b/arm4/arm5): p=.25. Thus place cell distribution in any of the arms was not significant compared to the rest across any of the tracks.

Further, when the above RUN distribution of both CA1 and CA2 place cells combined were collated according to novel and familiar areas on track on day 2 and 3, it was observed that on day 2, most place cells fired for novel part of the track (RUN2new) and that this count was similar to RUN1 place cell number. On the other hand, place cell number on RUN2fam decreased to almost half the original place cell number from RUN1. For day3, I expected that as seen on day2 (RUN2new>RUN2fam), maximum number of place cells would continue to fire for novel portion of the track; but the reverse was actually seen (RUN3fam>RUN3new). Most cells fired for familiar part of the track on day3 instead, for which the cell count dropped to almost half from that of the previous day (RUN2new>RUN3new). Also, the number of place cells firing for familiar portion of track on day 3 returned to similar counts as that on day 1, as opposed to day 2 where they had dropped to half their original count (RUN1/RUN2fam/RUN3fam) (fig3C).

Further, when the above said RUN distribution was divided into CA1 and CA2 ensemble populations separately, it was observed that both ensembles showed a very similar trend in their cell counts across RUN1, RUN2fam, RUN2new, RUN3fam and RUN3new. Both populations had higher cell counts for RUN2new as compared to RUN2fam, which in turn had comparable counts to RUN1. On day3, however, the reverse trend was observed and RUN3fam had a higher cell count than RUN3new, which in turn had a comparable count to RUN1. Thus, comparisons for place cell distributions across RUN2 and RUN3 for familiar and novel track parts seem to flip and be the opposite of one another (fig3D). (1-tailed, paired T-Test-CA1: RUN2fam/RUN2new: p=.074, RUN3fam/RUN3new: p=.007; RUN2fam/RUN3fam: p=.017, RUN2new/RUN3new:p=.056; CA2: RUN2am/RUN2new:p=.038, RUN3fam/RUN3new:p=.006; RUN2fam/RUN3fam:p=.09, RUN2new/RUN3new:p=.02).

The above myriad of comparisons give a clear picture of spatial area representation by place cell counts of CA1 and CA2 ensembles, both across and within days, as the environment is modified each day. As familiarity of the track increases from 0% on day1 to 50% on day2 and 67% on day3 (with 2/3rds of the track being relatively familiar now), it was correspondingly observed that 0% of place cells fired for familiarity on day 1 (RUN1), 35% of place cells fired for familiar track on day 2 (RUN2fam) and 70% on day 3 (combining RUN3fam, arm 2b, and arm 3a). Similarly, as novelty of the track decreased from 100% novelty on day1, 50% on day2 and 33% on day3, the corresponding % of place cells firing for novel track portion also decreased from 100% on day 1 (RUN1), to 65% on day 2 (RUN2new) to 30% on day 3 (RUN3new). Therefore, while for day 1 and day3, the correlation between the distribution of familiarity and novelty on track closely reflected the place cell distributions of both CA1 and CA2, but the same was not observed for day 2. There was a skewed representation of place cell distribution on day2, with respect to both familiarity and novelty. While the track on day 2 comprises of 50% familiar (arms 1, 2a,4b) and 50% novel track (arms 2b,3,4a), the place cell ensemble distribution for both cell populations was 35% for familiar track and 65% for novel track. This difference in place cell distribution between day 2 and 3 might indicate that when novelty is introduced in an environment for the first time (early detection), the neuronal ensemble response of both CA1 and CA2 is an increase in place cell number firing for that particular novel region, but not on subsequent detections (whenever novelty is introduced next) where both ensembles choose to redistribute its place cells in accordance with relative familiarity and novelty in the environment (fig3E).

### Place cell distribution across tracks

Place cell distributions were compared across all arms of all 3 tracks on all days, i.e. 3 arms on day1, 6 arms on day2 and 8 arms on day3 (figure1A). This was done to eliminate the possibility of attributing any of the above results to a preferential arm of any track by chance or for the reward area. It was observed that this was not on the case on either day 2 or 3. On day2, all 3 arms of the novel track showed an increase in number of place cells firing for it when compared to 3 arms of the familiar track (RUN2fam v/s RUN2new). Similarly, the result seen on day 3 was due to an increase in place cells firing for all 3arms of familiar track when compared with all 3 arms of novel track (RUN3fam v/s RUN3new). The place cell distribution across the particular arms connected to the reward area on any of the days also did not show a preferential increase/decrease compared to any other arm of the tracks. Thus, the place cells were pretty evenly distributed across all arms of novel tracks and familiar tracks, and were not influenced by either a particular arm of any track or the position of reward area on any day. (Kruskal Wallis test: RUN1 arms place cell distribution (arm1/arm2/arm4): p=.9, RUN2fam arms (arm1/arm2a/arm4b): p=.06, RUN2new arms (arm2b/arm3/arm4a): p=.67; RUN3fam arms (arm1/arm2a/arm6): p=.058, RUN3new arms (arm3b/arm4/arm5): p=.25) (fig3F-table)

Introduction of novel /added arms to an already pre-existing environment does result in higher number of place cells firing for that space, (Frank *et al*,2004), but as observed in the above results, this phenomenon occurs only during early detection (first novelty introduction) and not during subsequent detections of novelty. Nonetheless, it is imperative that no matter how many times novelty is introduced in an environment, the animal recognizes this addition/modification and behaves accordingly. Thus, to explore what the neuronal ensemble response might be during subsequent novelty detections (if not an increase in place cell count), differences (if any) in other firing properties of these pyramidal neurons were studied.

### Average firing rate of place cells

The average firing rate of all place cells from both cell populations were calculated for each arm of all tracks, across all days (see supplementary-figure2). It was observed that on day1, the arm with place cells having the highest average firing rate was arm2; while on day2, the highest firing rate was from place cells belonging to arm2b (the 1^st^ arm of Run2new) and on day3, were from arm4 (middle arm of Run3new) (fig4A). Even when observed for CA1 and CA2 populations separately, the same was observed (arm 3a and arm5 for CA2 had no place cells therefore have the value 0) (fig4B). This seems to suggest the possibility that place cells firing for novel part of the track have a higher firing rate than those firing for the familiar part.

Further, the firing rate of place cells on common arms from RUN1 (arm1,2,4) were compared with RUN2fam ( arm1,2a,4b) and RUN3fam (arm1,2a,6) correspondingly and it was seen that across all 3 arms, the firing rate decreased from day 1 to 2 to 3 (Kruskal Wallis test: RUN1/RUN2fam/RUN3fam p<.0001). The same was observed for CA1 and CA2 populations separately as well (Kruskal Wallis test: RUN1/RUN2fam/RUN3fam-CA1 :p<.0001; CA2:p=.0003) (fig4C). Further, the average firing rate on each of the 3 arms of the common tracks of RUN1, RUN2fam and RUN3fam were compared individually across days and it was observed that the firing rate decreased each day across each of the individual arms as well (Kruskal Wallis test: RUN1/RUN2fam/RUN3fam: arm1: p=.0002, arm2: p=.001, arm4: p<.001). The same was observed for CA1 place cells separately as well (Kruskal Wallis test: CA1: RUN1/RUN2fam/RUN3fam: arm1: p=.0025, arm2: p=.0008, arm4: p=.0001), but not for CA2. Although the same trend in each of the arms was observed in CA2 as well, it wasn’t statistically significant, probably due to less number of cells. Therefore, as a novel spatial area becomes more and more familiar, the average firing rate of place cells firing for that space decreases. Next, within day comparisons of firing rates were made between CA1 and CA2 place cells on RUN2fam and RUN2new and higher firing rates were found for place cells firing on RUN2new (fig4D), although not statistically significant. In contrast, when the same comparison was extended to RUN3, the difference between firing rates of RUN3fam and RUN3new were more pronounced and clear across all 3 arms RUN3new>RUN3fam (Mann Whitney test-RUN3fam/RUN3new:p<.0001). The same was observed for CA1 and CA2 separately as well (Mann Whitney test-RUN3fam/RUN3new: CA1:p=.0001; CA2:p.0219) (fig4E). (Note: arms 1,2,3, indicated in figure 4d and 4e do not correspond to actual arms 1 2 3 on track but are used for numbering the 3 arms of RUN2fam, RUN2new, RUN3fam and RUN3new. Also, arm3 for CA2 corresponds to arm5 on track (last arm of RUN3new), for which there were no place cells, hence only 2 arms are shown in figure). Furthermore, since RUN1 was completely novel on day1, the average firing rates were compared for RUN1/RUN2new/RUN3new for all cells together as well as CA1 and CA2 place cells separately, and it was observed that all 3 had similar average firing rates, thereby not showing a statistically significant test; indicating that average firing rates of novel place cells are similar to one another. (RUN1/RUN2new/RUN3new: Kruskal Wallis test: all place cells: p=.107; CA1 cells: p=.259; CA2 cells: p=.250).

**FIGURE 4:**
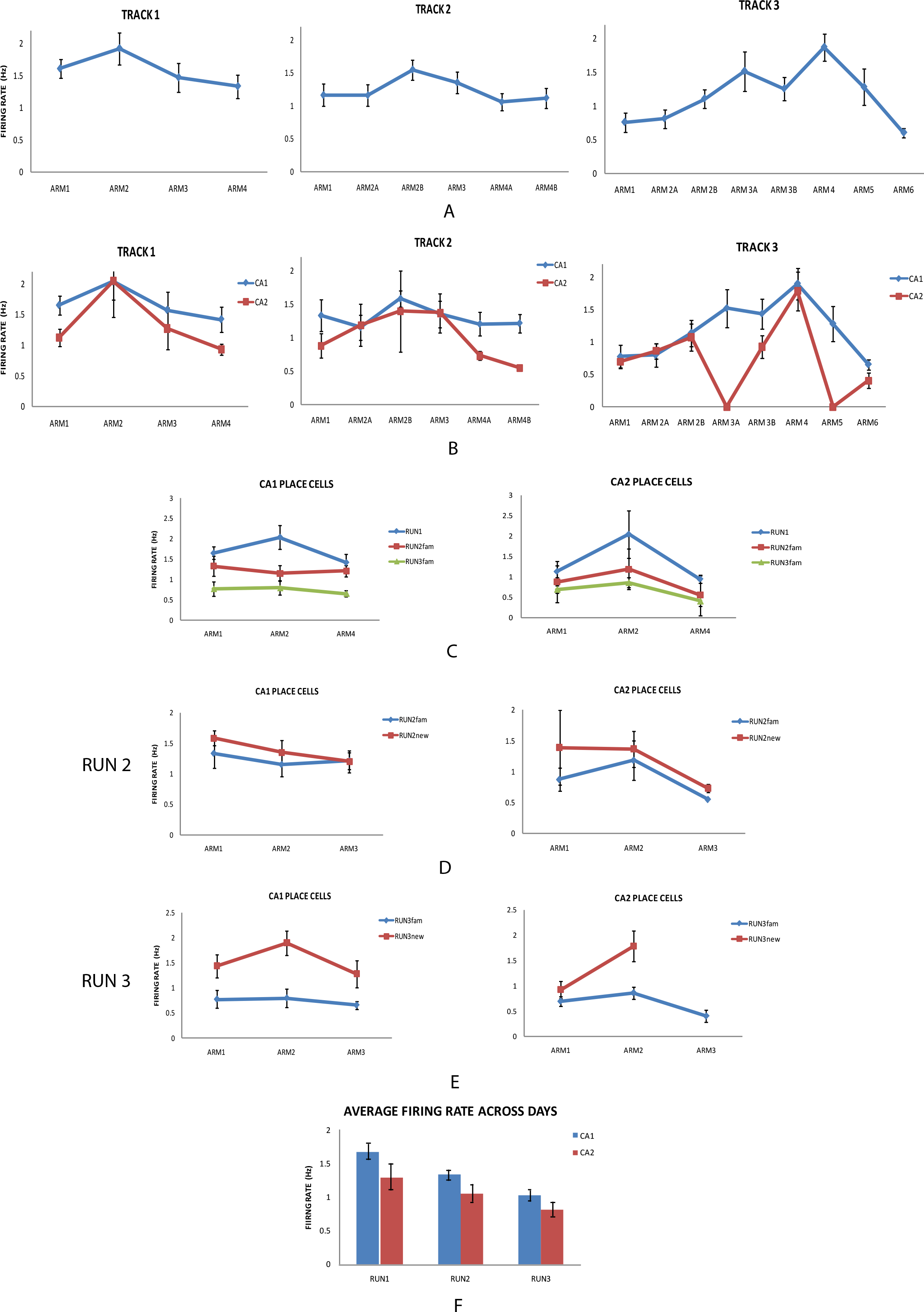
a) Average firing rate of all place cells (CA1+CA2) across all arms of all tracks each day. The arm with highest average firing rate on day 1 was arm2; on day 2 was arm2b (the 1^st^ arm of novel track RUN2new) and on day 3 was arm 4(2^nd^ arm of novel track RUN3new). b) The same trend was seen when the average firing rates of CA1 place cells and CA2 place cells were compared separately. c) Average firing rates across common arms of all tracks: RUN1/RUN2fam/RUN3fam comparison for individual arms across days: as the novel area of RUN1 became more and more familiar on day2 (RUN2fam) and day3 (RUN3fam), the average firing rates of place cells decreased each day.| Kruskal Wallis test :All cells: RUN1/RUN2fam/RUN3fam: p<.0001, CA1cells :p<.0001; CA2 cells:p=.0003. This trend was observed for within individual arms of the track as well (arm1 of RUN1 v/s arm1 of RUN2fam v/s arm1 of RUN3fam; arm2 of RUN1 v/s arm2a of RUN2fam v/s arm2a of RUN3fam; arm4 of RUN1 v/s arm4b of RUN2fam v/s arm6 of RUN3fam) for all cells combined as well as for CA1 cells separately. Kruskal Wallis test: All cells: arm1: p=.0002, arm2: p=.001, arm4: p<.001; CA1 cells: arm1: p=.0025, arm2: p=.0008, arm4: p=.0001. d) Within day comparison for RUN2: average firing rates for both CA1 and CA2 place cells firing in 3 arms of RUN2new were higher than the 3 arms of RUN2fam, but not statistically significant. e) Within day comparison for RUN3: average firing rates for both CA1 and CA2 place cells firing in 3 arms of RUN3new were significantly higher than the 3 arms of RUN3fam. Mann Whitney test for RUN3fam/RUN3new: All cells: p<.0001; CA1cells :p=.0001; CA2 cells :p=.0219. (Note: arms 1,2,3, indicated in figure 4d and 4e do not correspond to actual arms 1, 2, 3 on track but are used for numbering the 3 arms of RUN2fam, RUN2new, RUN3fam and RUN3new. Also, in fig 4e, arm3 for CA2 corresponds to arm5 on track (last arm of RUN3new), for which there were no place cells, hence only 2 arms are shown in figure). f) Average firing rates of CA1 and CA2 place cells across RUN1, RUN2 and RUN3. For both populations, the average firing rates decreased as the overall familiarity of the environment increased (0% familiarity on day1, 50% familiarity on day 2 and 67% familiarity on day 3). Kruskal wallis test for RUN1 v/s RUN2 v/s RUN3: CA1 cells: p<.0001; CA2 cells: p=.0442. Bars represent standard error of mean for each of the figures.

Finally, average firing rates for CA1 and CA2 were calculated across RUN1, RUN2 and RUN3 and it was observed that they decreased across days (fig4F), further cementing the previous observation that as a completely novel area gets more and more familiar (0% familiar on day1, 50% familiar on day2 and 67% familiar on day3 as stated above), the firing rates of place cells coding for that area decrease (Kruskal wallis test: RUN1/RUN2/RUN3: CA1:p<.0001; CA2:p=.0442). This distinction was also observed within the same day, for day 2 and 3, where both cell populations showed a higher average firing rate on RUN2new and RUN3new, compared to RUN2fam and RUN3fam respectively (fig4D, 4E). This indicates that the animal was able to distinguish between relative novelty and familiarity even when acquiring spatial episodic memories of a given environment in real time. It was also observed that the average firing rates of CA1 place cells were higher than those of CA2 place cells across all days.

### Pairwise cross correlations between novel and familiar place cell pairs

Previous studies have reported that CA1 cell pairs with overlapping place fields show a higher correlation and co-ordinated activity during high frequency events (HFE) in novel environments and decrease with increasing familiarity (Cheng & Frank, 2008). Therefore, I looked into pairwise cross correlation differences between novel cell pairs and familiar cell pairs on all tracks, across and within days. Those cell pairs were chosen that had overlapping place fields (peak distance<15cms) and were recorded on different tetrodes (Wilson & McNaughton, 1994; Skaggs & McNaughton, 1996). These cell pairs included CA1-CA1 pairs, CA1-CA2 pairs and CA2-CA2 pairs. Since CA2-CA2 pairs were relatively fewer in number, comparisons were limited to all cell pairs taken together, CA1-CA1 and CA1-CA2 pairs (see supplementary fig2-table). Pairwise cross correlations from all cell pairs from each arm of all tracks were compared, as previously done for average firing rates. It was observed that on day2, the highest correlation was between cell pairs belonging to arm 2b (the first arm of Run2new) and on arm 5, followed by arm 4 (2^nd^ and 3^rd^ arms of RUN3new) on day3. This is the same observational trend as seen when comparing average firing rates previously(fig5A). The same comparison was done for CA1-CA1 pairs and CA1-CA2 pairs separately as well: peak cross correlations were seen for cell pairs belonging to arms 4a (CA1-CA1 pairs) and arm2b (CA1-CA2 pairs) on day 2 (both arms belonging to RUN2new); and on arms 3b and 5 (CA1-CA1 pairs), arms 3b and 4 (CA1-CA2 pairs) on day 3 (both arms belonging to RUN3new) (fig5B).

Next, pairwise cross correlations between all cell pairs were compared across RUN1, RUN2fam and RUN3fam, and it was observed that they decreased across days (Kruskal Wallis test: RUN1/RUN2fam/RUN3fam: p<.0001). There was no correlation found for within arm comparison (arms1,2,4) for the same, as was found for average firing rates; therefore all arms were taken together and compared across days. The same was observed for CA1-CA1 and CA1-CA2 cell pairs separately as well (Kruskal Wallis test: RUN1/RUN2fam/RUN3fam: CA1-CA1 pairs: p<.001; CA1-CA2 pairs: p=.0009) (fig5C). For within day comparison on day 2 and 3, it was observed that correlations between pairs on RUN2new were higher than RUN2fam and for RUN3new than RUN3fam. The same was observed for CA1-CA1 and CA1-CA2 pairs separately as well. Although, as observed for average firing rates, the differences were more pronounced on day 3 than day2. (Mann Whitney test: RUN3fam/RUN3new-all pairs: p.0312; CA1-CA1 pairs: p<.001) (fig5D and 5E). (Note: as for average firing rates, arms1,2,3 mentioned in fig 5d and 5e do not denote actual arms1,2,3 of any track but are only used for numbering purposes. Also, in fig 5e, arm3 for CA2 corresponds to arm5 on track (last arm of RUN3new), for which there were no place cells, hence only 2 arms are shown in the figure and the statistical test could not be conclusive). Furthermore, the cross correlations between cell pairs on novel areas from all 3 days were compared as well (RUN1/RUN2new/RUN3new) and there wasn’t any significant difference found, similar to the trend observed previously for average firing rates as well; indicating that pairwise cross correlations between novel cell pairs are also similar in values. (RUN1/RUN2new/RUN3new: Kruskal Wallis test: all pairs: p=.088; CA1-CA1 pairs: p=.485).

**FIGURE 5:**
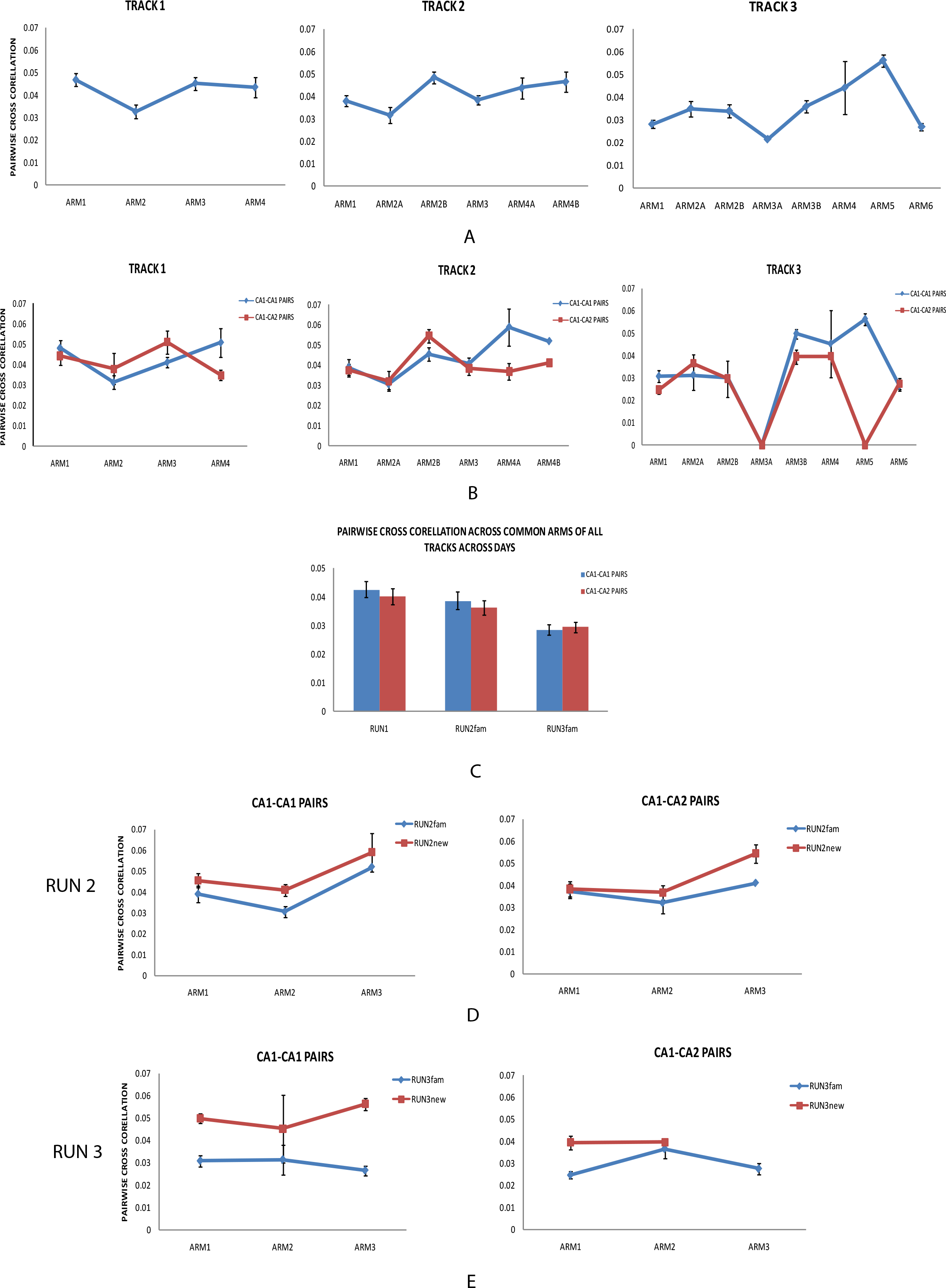
a) Pairwise cross correlations of CA1 and CA2 cell pairs (CA1-CA1, CA1-CA2, CA2-CA2) across all arms of all tracks each day. The arm with highest average firing rate on day 2 was arm2b (the 1st arm of novel track RUN2new) and on day 3 was arm5 followed by arm 4(1st 2 arms of novel track RUN3new). b) Similar trend was seen when Pairwise cross correlations were seen independently for CA1-CA1 place cells pairs and CA1-CA2 place cell pairs. c) Pairwise cross correlation across common arms of all tracks: RUN1/RUN2fam/RUN3fam. The correlations between cell pairs decreased each RUN with increasing familiarity of the track. Kruskal wallis test for RUN1 v/s RUN2fam v/s RUN3fam: All cell pairs :p<.0001. The same trend was seen individually for CA1-CA1 and CA1-CA2 cell pairs. Kruskal wallis test for RUN1 v/s RUN2fam v/s RUN3fam: CA1-CA1 cell pairs: p<.001; CA1-CA2 cell pairs: p=.0009 d) Within day comparison for RUN2: pairwise cross correlations for CA1-CA1 and CA1-CA2 cell pairs, firing in 3 arms of RUN2new were higher than 3 arms of RUN2fam, but not statistically significant. e) Within day comparison for RUN3: pairwise cross correlations for CA1-CA1 and CA1-CA2 cell pairs firing in 3 arms of RUN3new were significantly higher than 3 arms of RUN3fam. Mann Whitney test for RUN3fam v/s RUN3new :CA1-CA1 cell pairs : p<.001., CA1-CA2 cell pairs: p=.0312 (Note: arms 1,2,3, indicated in figure 5d and 5e do not correspond to actual arms 1,2, 3 on track but are used for numbering the 3 arms of RUN2fam, RUN2new, RUN3fam and RUN3new. Also, in fig 5e, arm3 for CA2 corresponds to arm5 on track (last arm of RUN3new), for which there were no place cells, hence only 2 arms are shown in figure). Bars represent standard error of mean for each of the figures.

### Spatial novelty consolidation during SWRs replays in sleep

While both CA1 and CA2 showed the same firing responses for both early spatial novelty detection and subsequent novelty detection, I further wanted to investigate for differences (if any) in spatial consolidation mechanisms between the two neuronal ensembles. Therefore I analyzed spiking activity within the confines of SWRs, isolated from nREM sleep, from each day’s post run. A total of 203 such significant sequences were detected on day1, 461 on day 2 and 314 on day 3. These sequences were further divided into 2 categories: CA1 sequences and CA1+CA2 sequences (since I could not find any sequences that only had spikes from CA2 cells in any rat on any day, and the probability of doing so is anyway extremely low owing to the diminutive physical area of CA2 as compared to CA1). The total number of sequences generated by both CA1 and CA2 place cells were highest for day2, and not day 3, despite the fact that the biggest spatial area traversed by the rat was on that day. (fig6A)

**FIGURE 6:**
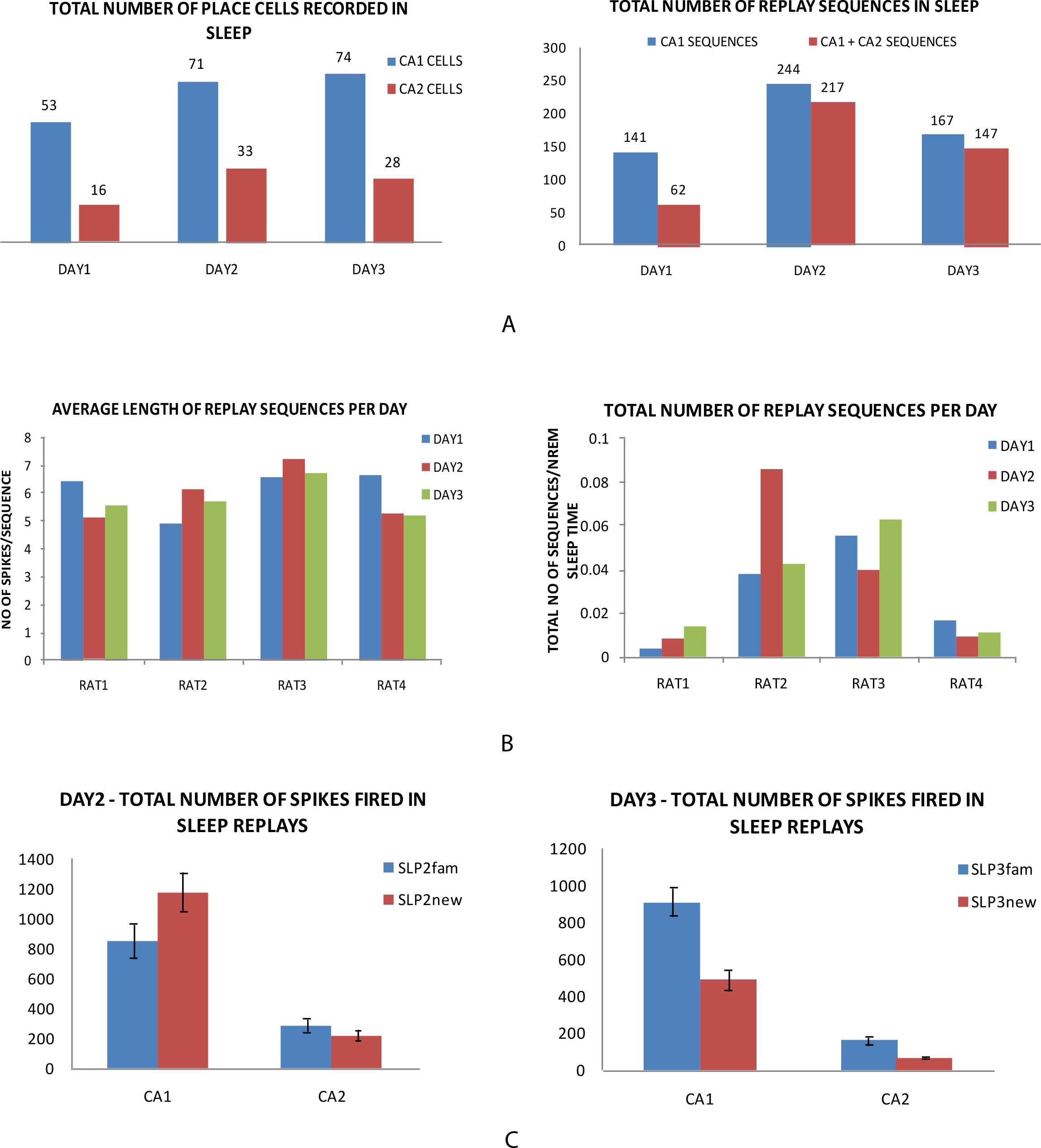
a) Total number of place cells recorded in sleep for CA1 and CA2 place cells. Total number of CA1 and CA1+CA2 replay sequences recorded in sleep each day. The highest number of sleep sequences were recorded on day2, despite a longer track length on day3. b) Average length of replay sequences across all days for all rats: As the track length increased, there was no consequential trend seen in length of replay sequences recorded in sleep. Total number of replay sequences recorded in sleep: there was no effect of increasing track length each day on the number of sequences recorded each day. c) Total number of spikes fired in sleep replays from CA1fam, CA1new, CA2fam and CA2new on day2 and 3. While for CA1, there was a coherent response observed each day in SLP for that particular day’s RUN, the same was not observed on CA2. CA1 spikes: 1 tail, paired T test: SLP2fam v/s SLP2new: p=.011; SLP3fam v/s SLP3new: p=.034. Bars represent standard error of mean.

Since the increase in the length of the track is the only change in the environment each day of the experiment, I wanted to check if this influences the length of replay sequences during SWRs in any way. As we go from day 1 to day 3, the track length almost doubles in size (72 spatial bins to 142 bins), therefore it was imperative to see how an ever-changing (length increase) environment would be consolidated in sleep. The average length of replay sequences were calculated and compared across days to see if they varied as a function of increasing length of the track. There was no such trend observed. Next, I checked if the total number of replay sequences recorded each day in sleep varied each day as the track length increased. Again, no trend of increase/decrease was observed across days for any of the rats (fig6B). Even when the total number of CA1 sequences and CA1+CA2 sequences were looked at separately, there was no correlation found with track length.

Next, the individual spikes firing in each of the significant replay sequences were classified further based on their peak firing positions on RUN for that particular day, and divided further into SLP1 (corresponding to RUN1), SLP2fam, SLP2new (corresponding to RUN2fam and RUN2new from day2) and SLP3fam and SLP3new (corresponding to RUN3fam and RUN3new from day3). Cumulative spikes fired in sleep sequences from all place cells belonging to either familiar or novel arms of the track on both day 2 and 3 were compared within the corresponding days. Each of the spikes firing within each sleep SWR, were classified as 1.CA1fam 2.CA1new 3.CA2fam and 4.CA2new for days 2 and 3. For day1, spikes were identified on basis of whether they belonged to CA1 or CA2. All the spikes belonging to one category were summed across and their cumulative totals were compared.

For within day comparison on day2, when CA1fam and CA1new were compared across all rats and it was observed that more spikes from place cells belonging to the novel track fired in SWR during sleep compared to their familiar counterparts. (1 tail, paired T test: SLP2fam/SLP2new: p=.011). This was the same observation in RUN2 (RUN2new>Run2fam). Similarly, for day 3, it was observed that a higher number of spikes fired in sleep from familiar part of the track (SLP3fam) across all rats when compared to novel part of the track (SLP3new) (1 tail, paired T test: SLP3fam/SLP3new: p=.034). This was again in sync with RUN3 (RUN3fam>RUN3new). Thus, for CA1, on both day 2 and 3, whichever part of the track in RUN had higher number of place cells firing for it, had a higher number of spikes firing in sleep from those place cells belonging to that part of the track. Therefore, I observed a ‘coherent response’ from CA1 place cells in RUN and SLEEP firing responses for a particular day (fig6C, supplementary-fig1).

Conversely, the same was not observed for CA2 spikes in sleep. For both day 2 and 3 when the cumulative total for SLP2fam v/s SLP2new and SLP3fam v/s SLP3new were compared, for some rats, more spikes from novel track fired more in sleep while for others, spikes from place cells belonging to familiar track fired more (fig6C, supplementary-fig1). Thus, there was no clear, conclusive trend seen for CA2 cells firing in sleep replays, and they showed no preference for any particular part of the track: familiar or novel, irrespective of how many place cells fired for each of those parts in RUN. (1 tail, paired T test: SLP2fam/SLP2new: p=.14; SLP3fam/SLP3new: p=.23). This is not only in contrast to CA1’s spiking response in SLEEP but, more importantly, to CA2’s own firing response in RUN on both day2 and 3 (which was exactly similar to CA1’s RUN on both days).

Therefore, even though CA1 and CA2 showed the same response and preference to a particular track part in RUN on both days, the ‘coherent response’ observed between SLEEP and RUN in CA1 was not observed in CA2. Thus, a stark difference in spatial memory consolidation mechanisms between CA1 and CA2 was observed.

Finally, spatial (track) coverage in RUN and SLEEP sessions across all days was plotted from CA1 and CA2 place cells combined for each track. If SWRs in sleep are a form of memory consolidation (Wilson & McNaughton, 1994; Kudrimoti *et al*., 1999; Nádasdy *et al*., 1999; Lee & Wilson, 2002), it is imperative to see how a completely novel environment (day1) is represented by CA1 and CA2 in sleep and on subsequent days, when it becomes increasingly familiar (days 2 and 3), while the track also simultaneously changes form, with respect to shape and length. All place cells (CA1+CA2) that fired on track in RUN sessions (RUN1, RUN2fam, RUN2new, Run3fam, RUN3new) for that particular day were plotted for that particular track, in accordance with their peak firing position. Since the place cells corresponding to reward area and any spatial area outside of RUN were eliminated for analysis, some gaps may be observed in the plotting. (fig7-RUNl, RUN2, RUN3). Out of those place cells, cells that fired in replays corresponding to SLP1, SLP2fam, SLP2new, SLP3fam and SLP3new were plotted for all days, respectively (fig7-SLP1, SLP2, SLP3). For plotting purposes, only the 1^st^ spike from each cell that fired multiple times in a replay sequence was taken per sequence. When all cumulative replay sequences were plotted together, I observed that on day 2, place cells corresponding to SLP2new fired more than SLP2fam, while the reverse was seen on day 3 (SLP3fam>SLP3new). This is in congruence with more place cells (CA1 and CA2) firing in RUN2new when compared to RUN2fam and RUN3fam when compared to RUN3new. This is also in accordance with CA1 cell firing responses in sleep, which in itself was congruent with its own RUN responses on both day 2 and 3 (supplementary-fig1)

**FIGURE 7:**
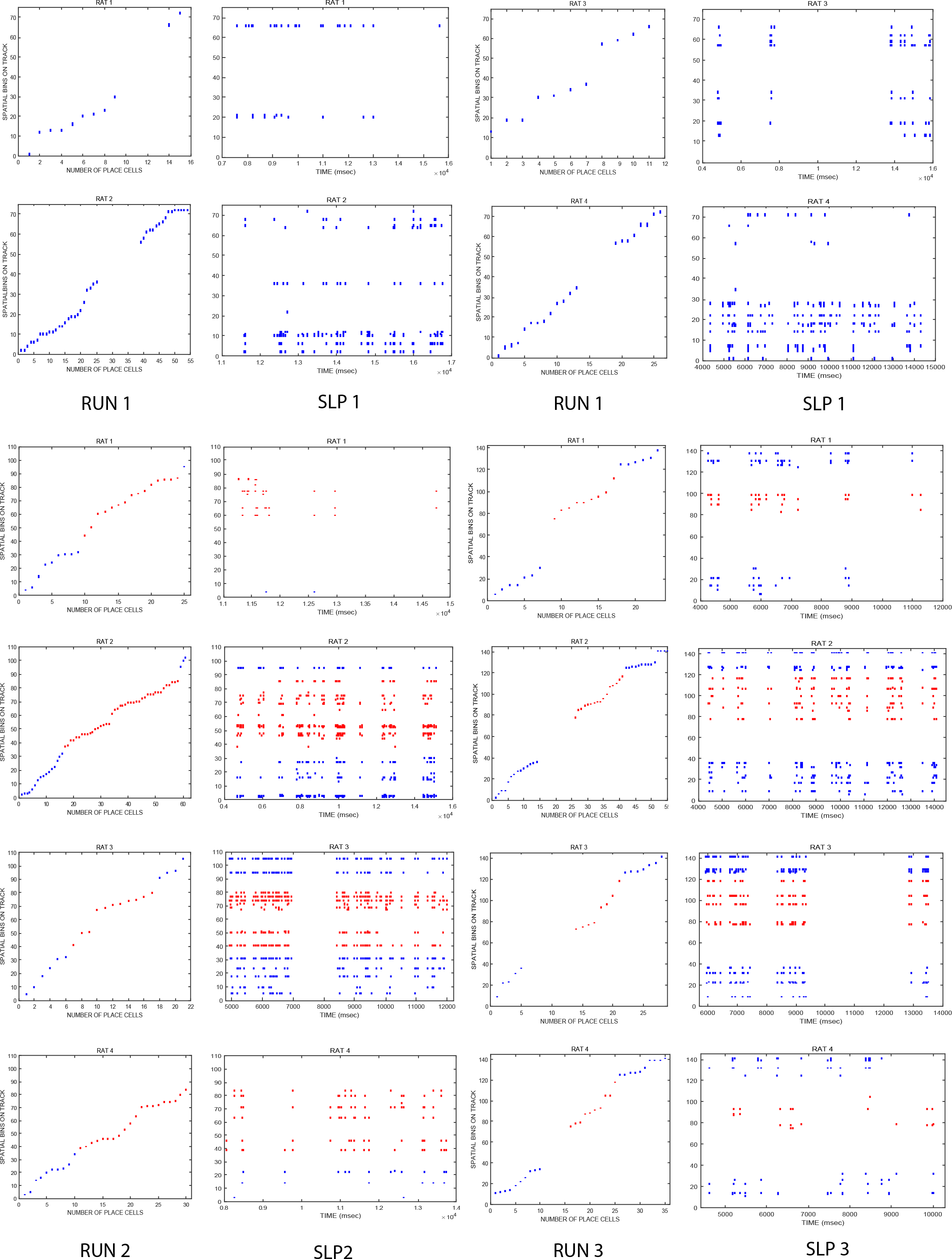
a) Representation of all tracks in RUN and SLP for each rat: each rat’s recorded place cells from CA1 and CA2 are plotted for RUN1,RUN2,RUN3 separately in the left panel. Each rat’s replay sequences from the corresponding SLP session are plotted as SLP1,SLP2,SLP3 on the right panel. Red denotes spikes fired from place cells firing for novelty in RUN2new/SLP2new and RUN3new/SLP3new while blue denotes spikes from place cells that fire for familiarity in RUN1/SLP1, RUN2fam/SLP2fam and RUN3fam/SLP3fam. On day2, there are significantly more number of place cells for RUN2new firing in sleep replays (SLP2new) as compared to RUN2fam (SLP2fam). The opposite is seen on day3, where place cells from RUN3fam fire more than RUN3new in sleep (SLP3fam>SLP3new).

## Discussion

The aim of this study was to elucidate how hippocampal ensembles, particularly CA1 and CA2, dynamically evolve, modify and code for spatial familiarity and novelty in a given environment. When novelty is first introduced in a given environment, both ensembles devoted more number of place cells to encode for the novel area, as compared to the relatively familiar one, even though both spaces were similar in length, spatial structure and form, i.e. C-shaped tracks (RUN2fam/RUN2new). Additionally, place cells from both ensembles coding for the novel space had a relatively higher average firing rate and pairwise cross correlation compared to place cells firing for familiar locations as well, though not achieving statistical significance. Thus, the very first introduction of novelty resulted in a disproportionate allocation of place cell distribution (65% place cells coding for 50% novel area, 35% place cells for 50% familiar area). Conversely, when novelty was introduced on the subsequent day (day 3), no such disparity in place cell distribution was seen in any ensemble (30% place cells coding for 33% novel space, 70% place cells for 67% familiar space). Nevertheless, the place cells coding for novel space did have a significantly higher average firing rate and pairwise cross correlation than their familiar counterparts (RUN3fam/RUN3new). In addition, as a novel space became more and more familiar over successive days, the average firing rate and pairwise cross correlation for cells coding for this area decreased with increasing familiarity (RUN1>RUN2fam>RUN3fam). These observations indicate how hippocampal ensembles continually evolve, update and code for novelty and familiarity in the same space, in real-time, as the rat traverses through the given environment, and how robustly it does so, even as relative familiarity and novelty is experienced in the same lap itself (days 2 and3). Thus it seems that even within laps, the brain is able to distinguish between the relative novelty and familiarity pretty well, and this ‘novelty coding’ is rooted in specific characteristics of the novel place cells that develop as the rat traverses that area for the first time.

The second part of the study focused on better understanding the assimilation and consolidation of such an ever-changing environment across days. If ‘replay’ of place cells occurring during SWRs in nREM sleep is how spatial memory consolidation takes place, then it is important to study the preferential activation of place cells (if any) that takes place during these SWRS and to which part of the track/area they belong to. Particularly, with respect to this paradigm, given that relative familiarity and novelty existed in the same closed loop track itself, it is essential to observe how different parts of the track are represented in sleep each day. For this aim, I counted and combined all spikes that fired during significant replays in sleep, from all place cells that fired from various parts of the track in prior behaviour for each day. For CA1, whichever part of the track had a higher number of place fields in RUN session for that particular day, spikes from those place fields fired more in subsequent sleep as well. This consistent/coherent response was observed on both days (RUN2/SLEEP2 and RUN3/SLEEP3). The same was not observed for CA2, which showed no clear preference for either part of the track in sleep replays on any of the days, despite having a clear preference in RUN2 and RUN3, in terms of number of place cells firing for a particular spatial area.

### Early and subsequent spatial novelty detection is not the same

In vivo data demonstrating how CA1 and CA2 interact and contribute to continual spatial novelty detection and its subsequent consolidation are sparse, with most studies focusing on either social or novel contextual changes in an environment (Wintzer *et al*., 2014; Alexander *et al*., 2016) or non-spatial aspects of memory (Mankin *et al*., 2015). Other studies have primarily focused on novelty detection and encoding in CA1 only, in an object-place recognition task (Larkin *et al*., 2014), place field plasticity (Frank, 2004) or temporal coding and episodic memory (Mankin *et al*., 2012). To my knowledge, this is the first time where the roles of both CA1 and CA2 have been simultaneously examined with respect to spatial novelty detection and its subsequent consolidation. I wanted to observe how an existing hippocampal place cell ensemble for a completely novel environment (day1) would reorganize to incorporate the increasing spatial RUN area across days (days 2 and3). I also wanted to observe how place cell redistribution would occur as many new place cells would come up for the extended part of the track each day, and many would continue to fire for the familiar portion of the track. This would include activation of previously ‘silent’ place cells as well as some active place cells remapping to fire for another part of the track. Given the fact that the same hippocampal ensemble is highly malleable and flexible in encoding various different environments, it is imperative to see how the same ensemble would modify/reorganize to parametric variation of geometry, i.e. length of track. Given the ‘nontopographical mapping’ of any environment by hippocampal place cells (O’Keefe, 1976; Wilson & McNaughton, 1993), this question becomes even more pertinent and of greater importance.

For both CA1 and CA2 ensembles, on day 2, more place cells fired for the added portion of the track compared to the existing part of the track, while the reverse was seen on day 3. This makes sense as it isn’t energy efficient or physically possible for a given hippocampal place cell ensemble to keep modifying its distribution of place cells every time a spatial area is added to an existing environment, nor can it afford to dedicate more and more place cells that fire for the novel area each time it is introduced in the environment, as it would result in a highly uneven and inadequately sparse distribution of place cells for the entire spatial area.

But it is essential that each time modifications such as elongation of a track etc are implemented in an otherwise stable environment (with respect to other spatial cues, reward area, beginning/end of lap etc), the animal acknowledges this change and recognizes the novel/added spatial area. This is probably the reason why even though on the 3^rd^ day, number of place cells that came up for the added/novel track were lower in number, they still had a significantly higher average firing rate and a higher pairwise cross correlation between them when compared to their familiar counterparts.

Therefore spatial novelty detection seems to be a complex, dynamic mnemonic process, characterized by different and distinct hippocampal pyramidal neuronal properties. While early/first time response to spatial novelty detection is characterized by a higher number of place cells firing for the novel area, subsequent novelty detections are expressed through other spatial firing properties and characteristics of novel place cells such as higher average firing rate and pairwise cross correlations. Thus, the biggest difference between the hippocampal neuronal responses to novelty on day 2 and day 3 is higher number of place cells firing for the novel area when compared to the familiar one on day 2 but not day 3. Additionally, average firing rates and pairwise cross correlations between place cells firing for novel areas on all days (RUN1/RUN2new/RUN3new) were comparable to one another, indicating that novelty detection is expressed through these specific attributes and characteristics of novel place cells. Other characteristics of such novel place cells, not examined in this study may also contribute to subsequent spatial novelty detection as well, and is a bright future prospect to explore further.

I report for the first time that not only CA1 but also CA2, a small area, 1-step upstream and very much capable of influencing CA1 also shows the same response for both early and subsequent spatial novelty detections. Previous studies have shown that both CA3 and CA1 place cells are less stable in novel v/s familiar environments (Leutgeb, 2004), whereby CA1 input is largely dominated by the powerful disyanptic pathway of EC-CA2-CA1, and is only later on taken over by the classic trisynaptic pathway, over days (post 24 hours) to stabilize CA1 place fields (Karlsson & Frank, 2008). Lesion studies of EC-CA1 pathway have impaired spatial coding (Brun *et al*., 2008), indicating that CA2 directly influences spatial acquisition and learning driven processes in CA1. CA2 is uniquely poised within the hippocampal circuitry, with strong unidirectional connections to CA1 (Chevaleyre & Siegelbaum, 2010) and bi-directional feed forward connections to CA3 (Kohara *et al*., 2014; Boehringer *et al*., 2017). While dorsal CA2 projections to ventral hippocampus are required for social recognition memory (Meira *et al*., 2018), the same projections (excitatory) to dorsal CA1 seem to be involved in maintaining sequential firing patterns and working memory in CA1 (MacDonald & Tonegawa, 2021). Further, changes in an environment such as shape (Mankin *et al*., 2015) affect CA2 place fields the least when compared to CA1 or CA3 place fields. On the other hand, their sensitivity to smaller contextual changes and local cues, suggests that CA2 activity is an indicator of novelty signal to its downstream areas, i.e. CA1. Additionally, CA2 also receives projections from novelty signalling areas such as supra-mammillary nucleus (Chen *et al*., 2020) and non dopaminergic inputs from ventral tegmental atea (VTA) (Lisman & Grace, 2005). Thus, during the development of novel place fields, the input to CA1 is primarily dominated by CA2, which in itself codes for a novelty signal, and hence may explain why both ensembles show the same characteristic response to both types of novelty detection (early and subsequent).

### Place cell representation of a dynamic environment in sleep replays differs in CA1 and CA2

For a successful navigation strategy, mere mapping of novel geometry and its encoding is not enough. Assimilation of novel stimulus with existing memory traces of the environment and remembrance of various paths and spatial cues (proximal and distant) is equally essential. Consolidation of spatial memory is dependent on the re-activation of place cells that fired during recent behaviour and this replay is the cornerstone of higher order representations, which are spatial as well as non-spatial in nature. Patterns of network activity that occur within sharp wave ripples during sleep/awake resting pauses are on a time compressed scale that reflect recent tasks and episodic behaviour (Kudrimoti *et al*., 1999; Lee & Wilson, 2002; Davidson *et al*., 2009). This study would not be complete without focusing on learning and memory consolidation mechanisms in CA1 and CA2 and understanding their contribution to these mnemonic processes. I focused on spatial memory consolidation, particularly during sharp wave ripples during nREM sleep, which has been well established to be playing a crucial role in these processes (Buzsáki, 1986; Maquet, 2001; Marshall & Born, 2007).

As the track changed geometry and introduced familiar and novel spatial areas across days, I focused on each track’s representation in that corresponding day’s sleep by CA1 and CA2 place cell firing during SWR-replays and how it would be consolidated each day. To the best of my knowledge, this is the first study to focus on spatial representation of a changing environment in sleep by the CA2 neural network. While a couple of studies have recently come up on studying activation of CA2 neurons in sleep, they have focused on either social memory (Oliva *et al*., 2020), its role during immobility (Kay *et al*., 2016) or its role in triggering SWRs (Oliva *et al*., 2016).

I found that contrary to my hypothesis that CA1 and CA2 place cells from novel/added area of the track would be represented more during SWRs in sleep, i.e. more spikes firing from novel place cells than place cells from familiar parts of the track, neither CA1 nor CA2 showed this ‘novelty preference.’ This may indicate that other areas of the hippocampal-parahippocampal network are actually involved in spatial novelty consolidation or that novelty consolidation mechanisms are not just limited to/reflected in replay activity during nREM sleep but rather are a result of much higher and complex network mechanism in the brain.

Nonetheless, I did find differences in CA1 and CA2 firing patterns in sleep; which was surprising, given that in RUN, both ensembles behaved the exact same way on all days, with respect to both early and subsequent spatial detection. While CA1 showed a ‘coherent response’ in SLEEP with that corresponding day’s RUN, CA2 did not. Thus, for CA1, whichever part of the track for a particular day had a higher number of place cells firing for it in behaviour, had a higher number of spikes firing in that day’s sleep replay as well (for day2 : RUN2new and SLEEP2new; for day3 : RUN3fam and SLEEP3fam). In contrast, for CA2, irrespective of whichever part of the track was represented more in behaviour (for day2: RUN2new; for day3: RUN3fam), there was no such clear preference in sleep. This seems to indicate that CA2 place cells fired from all parts of the track with similar/same probability in sleep replays, irrespective of its coding for familiarity or novelty in RUN/behaviour. This also seems to indicate a dichotomy for CA2 neural network ensemble, whereby a clear preference is shown by it for a particular track region during spatial novelty detection but is lost during consolidation (subsequent sleep).

Relatively minimal work on CA2 network dynamics and mechanisms with respect to various aspects of spatial learning and memory leaves a lot to desire when making interpretations or hypothesis of such results. The inconclusive trend observed for CA2 in sleep could also be attributed to less number of CA2 cells recorded, in comparison to CA1, and consequentially less number of spikes fired in sleep by those CA2 cells. Although, this did not hinder the consistent trend that was observed in CA2 RUN, with respect to place cell number, average firing rate or pairwise correlations. Therefore, while in RUN sessions across days, CA1 and CA2 show the same neuronal responses (which could be attributed to the independent influence of CA2 over CA1 via the disyanptic loop in place field development), in SLEEP sessions the case is not so. The differing spiking responses between the two heavily interconnected regions in sleep replays seems to indicate that the role of CA2 in sleep goes much beyond just directly influencing CA1 firing and instead has a much larger role to play in influencing and co-ordinating the overall hippocampal circuitry. CA2 receives a direct input from the supramammilary nucleus (Haglund *et al*., 1984; Berger *et al*., 2001) and has been shown to be the initiator of SWRs in awake immobile and sleep, acting as a trigger and initiator for sleep replays (Oliva *et al*., 2020). It also plays a role in coding for current location during immobility and sleep through brief periods of desynchronization in slow wave sleep (SWS) (Kay *et al*., 2016). Chemogenetic inactivation of CA2 during a novel spatial task resulted in poor replay quality and fidelity during subsequent sleep and a lower ripple power in CA1 (He *et al*., 2020). Further, the N units/ramping cells in CA2 may help in choosing and selecting which particular experience will be re activated during a subsequent sharp wave ripple in sleep (Kay *et al*., 2016; Stöber *et al*., 2020). More studies have indicated that the competitive, alternate and independent circuitries via CA3 (trisynaptic pathway) and CA2 (disynaptic pathway) to CA1 as well as excitatory projections from CA2 to CA3 (Kohara *et al*., 2014) modulate the flow of spatial information in the hippocampus depending on different behavioural states (sleep v/s awake). It has been proposed that CA2 drives sensory based representations in awake state while CA3 drives memory based representation in sleep (Middleton & McHugh, 2020) via adenosine; which allows CA3 to control sleep replay content. Therefore, CA3 takes over (via the classical trisynaptic loop) and influences CA1 firing in sleep replays more for memory consolidation of prior experience. CA3 itself has been recently shown to preferentially replay novel trajectories with higher fidelity in nREM sleep (Hwaun & Colgin, 2019). The influence of CA3 over CA1 could explain the coherent response seen in CA1 for both day2 and 3’s SLEEP, whereby CA3 specifically focuses on novel trajectories irrespective of RUN, whereas CA1 focuses on reflecting trajectories that had a higher number of place cells coding for it on that particular RUN. This ‘switch’ between dominance of influence over CA1 from CA2 in RUN/behaviour and CA3 in sleep could explain why CA1 and CA2 show the same response to spatial novelty detection but not in subsequent consolidation. This could be the first step towards better understanding what each of these hippocampal subfields independently contributes to during memory consolidation of space. Most replay studies have focused primarily on CA3 or CA1 (Buzsáki, 1986; Lee & Wilson, 2002; Csicsvari *et al*., 2007; Davidson *et al*., 2009), and more still have looked at fidelity of replays, whether in sleep or awake state, (Foster & Wilson, 2006; Karlsson & Frank, 2009) or on direction of replays (forward/reverse) (Diba & Buzsáki, 2007). I have instead chosen to focus on what the replays themselves constitute/ are comprised of, and what the representation from each place cell that fired from a particularly specific spatial area in prior behaviour might reflect as a spike fired in a temporally compressed manner during sleep. The relative preferences of firing of different trajectories during sleep replays with respect to familiarity and novelty on track by these subfields (CA3, CA2, CA1) can help in understanding memory consolidation of a dynamic environment better in the hippocampus.

In conclusion, the questions that I have aimed to ask in this study is through a completely novel paradigm specially designed to answer and shed light on some important characteristics of hippocampal neuronal ensembles that have been unexamined or overlooked. The paradigm is especially suited to better understand how place fields themselves develop in a novel environment (day 1) and how these ensembles redistribute their place cell ‘allocation’ subsequently when varying degrees of familiarity and novelty is introduced (days 2 and 3). This relative familiarity and novelty is experienced by the animal not only in varying degrees across days, but also within the same laps on days 2 and 3, making it a powerful tool to study the influence of spatial novelty and familiarity on hippocampal place cell development, coding and distribution, and subsequently for higher network mnemonic processes. To the best of my knowledge, this is the first study that has aimed to tease apart differences in spatial early detection and subsequent detection with respect to novelty. It is also the first to shed some light on spatial memory consolidation mechanisms in sleep replay by looking at what the replay events actually constitute, in terms of spatial area representation, and not just the fidelity of those replays. Finally, this study opens avenues where detection, encoding and consolidation mechanisms in CA1 and CA2 and be understood better and bigger, bolder questions may be asked through this paradigm to understand various higher-order mnemonic processes and hippocampal network interplay better.

### Ethics statement

All the procedures (animal care, surgical procedures and euthanasia) were performed in accordance with NIH guidelines and were approved by the Institutional Animal Ethics Committee (IAEC) of National Brain Research Centre at Manesar, Haryana, constituted by the Committee for the Purpose of Control and Supervision of Experiments on Animals (CPCSEA), Government of India.

### Data availability statement

For data and codes used in this manuscript, a detailed request may be sent to the corresponding author and as per institutional guidelines will be shared.

## Acknowledgements

The author would like to thank the academic supervisor Dr Arpan Banerjee (Scientist V/ Additional Professor, NBRC) for supervision and guidance in writing this manuscript; Mr. Indrajith Ramachandran, and Mr. Apoorv Sharma for their constructive feedback towards proofreading the manuscript; and Dr Collins Assisi (IISER, Pune) for his contribution in writing Matlab code for extracting reply sequences in sleep.

---

**SUPPLEMENTARY-FIG1.**
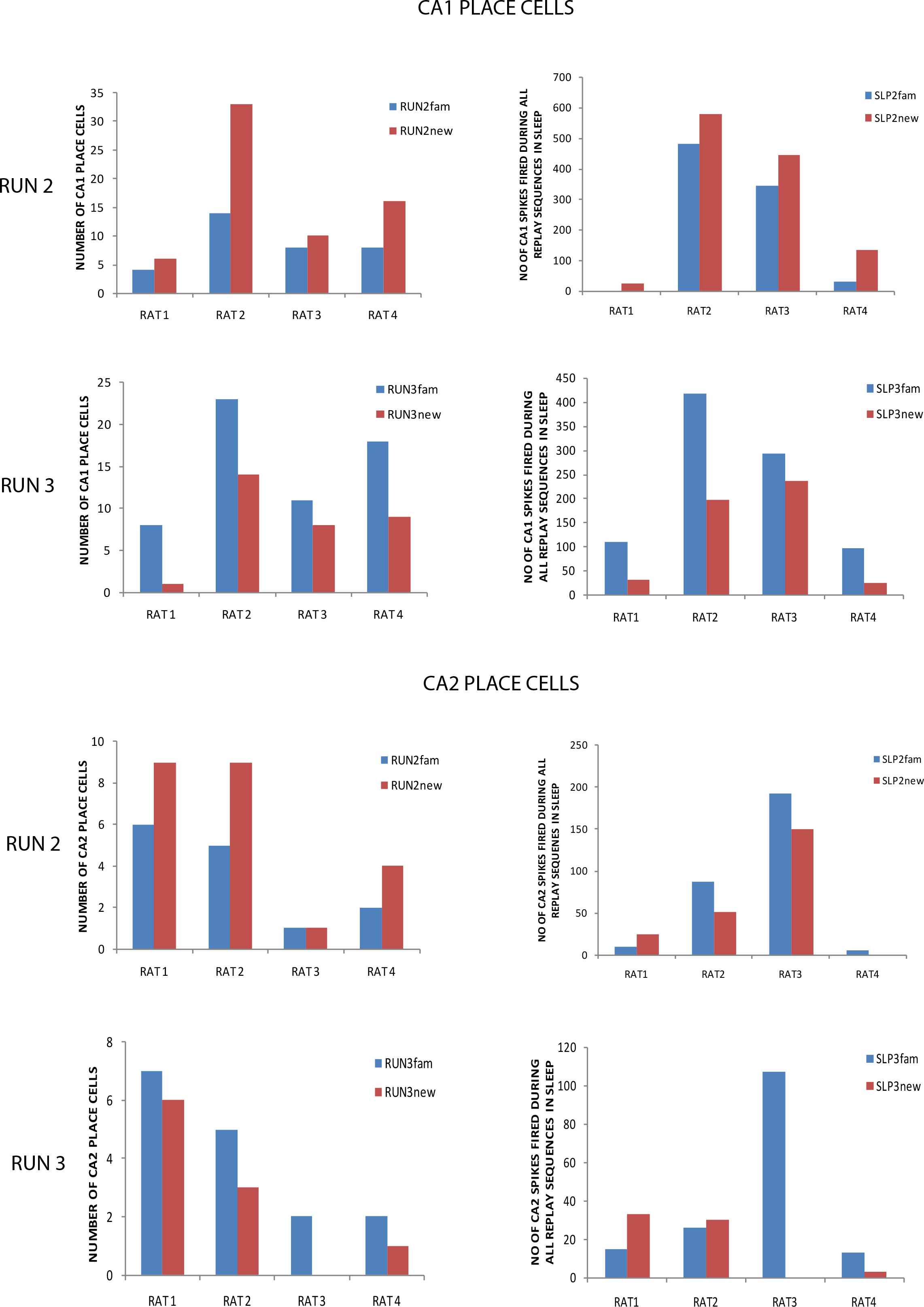
Hippocampal ensemble response in RUN and SLEEP for each rat: a) CA1 place cells: Day2: RUN2 and SLP2 show the same response as higher number of place cells fire for RUN2new than RUN2fam and correspondingly in SLP, spikes from place cells belonging to SLP2new fire more than spikes from SLP2fam during SWR-replays. Day3: similar to day2’s response, RUN3 and SLP3 show the same response. Higher number of place cells fire for RUN3fam than RUN3new and correspondingly in sleep spikes from place cells from SLP3fam fire more than SLP3new during SWR-replays. Thus, on both days, CA1 shows a ‘coherent response’ between a particular day’s RUN and SLP session. b) CA2 place cells: conversely, CA2 cells do not show the ‘coherent response’ on either day 2 or day3. While RUN2 and RUN3 responses mirror that of CA1’s RUN2 and RUN3 response (RUN2new>RUN2fam; RUN3fam>RUN3new), SLP responses (SLP2 and SLP3) do not show any conclusive trend. Thus, CA2 does not seem to show any preference for a particular part of the track as it did in its RUN responses (novel in RUN2 or familiar in RUN3).

**SUPPLEMENTARY-FIG2-TABLE.**
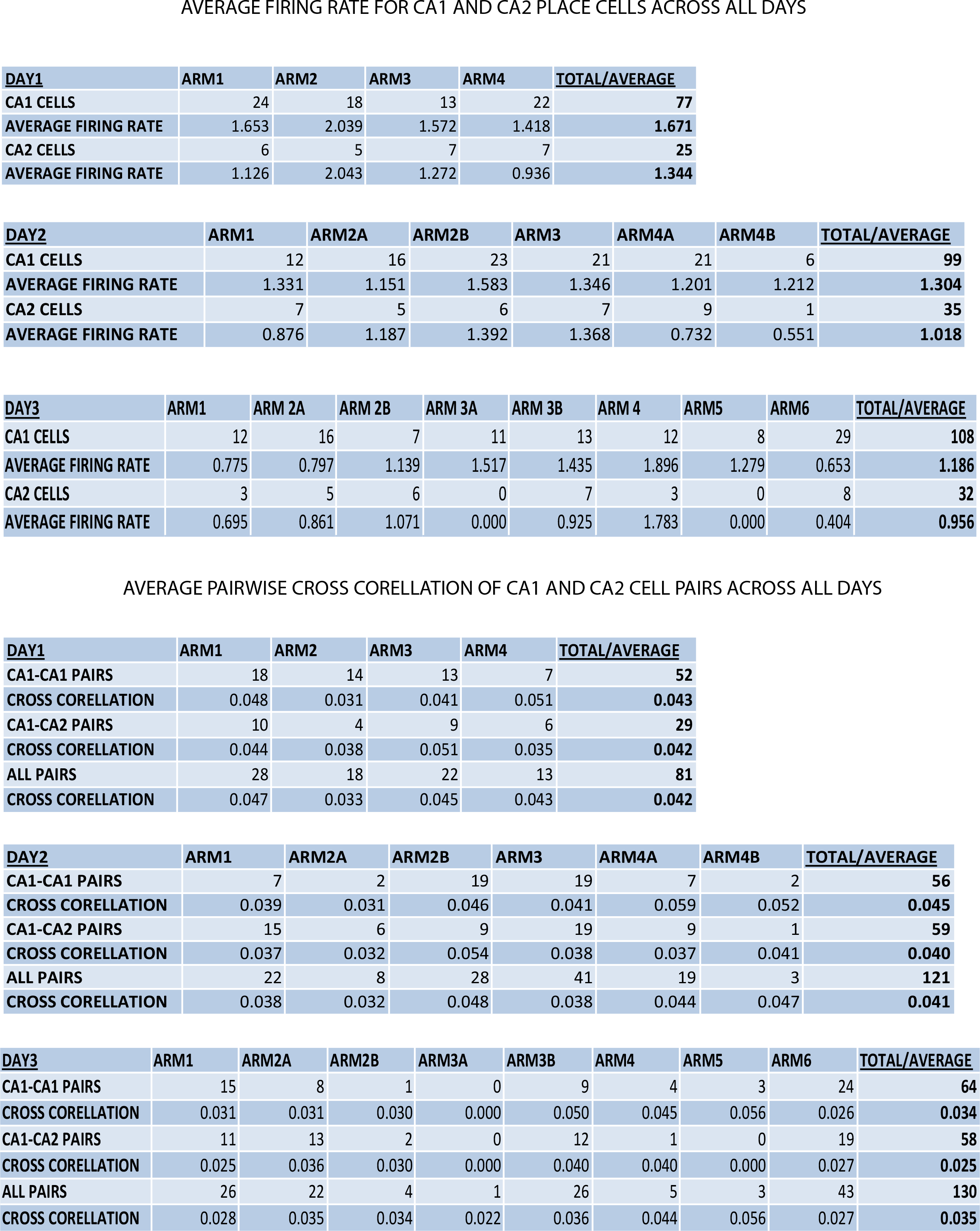
a) Total number of CA1 and CA2 place cells in each arm of all tracks across days and their corresponding average firing rates. a) Total number of CA1-CA1, CA1-CA2 and all (CA1-CA1,CA1-CA2,CA2-CA2) place cell pairs in each arm of all tracks across days and their corresponding pairwise cross correlation values.

